# On the energetic boundaries of trophic systems

**DOI:** 10.1101/2025.10.20.683547

**Authors:** Serguei Saavedra, Chengyi Long, Sonia Kéfi, Simon A. Levin, Pablo A. Marquet, Rudolf P. Rohr, Marco Tulio Angulo

## Abstract

Energy is the fundamental currency of life. Every organism requires a minimum intake to meet maintenance needs, and every trophic transfer dissipates part of the energy flow as heat. Classical theory has emphasized *lower* bounds on supply—the minimum resource level required for consumers to persist—yet the possible *upper* bounds and their role in limiting food-chain length remain unclear. We develop a thermodynamically grounded framework that reveals a triad of boundaries: (i) consumer thresholds, (ii) a basal protection limit that reframes the paradox of enrichment as a thermodynamic constraint, and (iii) environmental or physiological supply caps. We show that the intersection of these three boundaries determines the exact range of conditions under which a given ecosystem structure can occur, and that within this range trophic configurations arise as the unique solution that maximizes energy flow. As chain length increases, consumer thresholds rise while the basal protection limit falls, causing this range to contract and eventually disappear. Whether this limit is reached depends on the relative position of these two boundaries and the external supply cap. Their crossing defines an energetic ceiling: a thermodynamic bound on food-chain length beyond which no further level can be sustained. Our results show that feasibility conditions are directly shaped by thermodynamic laws, providing a mechanistic explanation for the emergence and limits of trophic systems.

## Introduction

About a century ago, Boltzmann, Lotka, and Schrödinger were among the first to recognize how life is in a continuous struggle to acquire enough energy to survive and reproduce (Boltzmann, 1905, Lotka, 1922, Schrödinger, 1944). At the level of individual organisms, this struggle has led evolution by natural selection to preserve developmental processes that economize energy. A central empirical pattern is the scaling between body size and metabolic demand, often summarized by Kleiber’s law (Kleiber, 1932, West et al., 1997), which posits that larger organisms tend to have lower mass-specific metabolic rates than smaller ones. This pattern has been interpreted as a consequence of constraints on energy supply and dissipation across transport networks, though recent work has emphasized that scaling exponents can vary across taxa and environments (Yeakel et al., 2018).

At larger levels of organization, populations form complex networks of energy supply and exchange (Odum and Barrett, 2005, Pielou, 2001). Ecosystems do not reproduce as units, yet filtering and feedback processes tend to favor systems with properties that enhance their robustness (Levin, 1999, 2005, Lewontin, 1969, Saavedra, 2024). This has motivated decades of work to understand how organismal energy requirements translate into ecosystem-level constraints (Brown et al., 2004, Enquist et al., 2003, Hatton et al., 2019, MacArthur, 1955). Classical ecological theory has long recognized energetic boundaries (Elton, 1927, Hutchinson, 1959, Lindeman, 1942, Schoener, 1989). Resource competition theory formalized the concept of a minimum requirement, or *R*^∗^, below which consumers cannot persist (Tilman, 1980). Matrix-based and structural approaches further emphasized how feasibility conditions can be expressed as inequalities identifying these lower bounds (Logofet, 1993, Saavedra et al., 2017). At the ecosystem scale, research highlighted how biodiversity and energy flows are constrained by such minima (Loreau, 2010). Consumer–resource models also revealed the paradox of enrichment, in which high nutrient input destabilizes basal populations (Rosenzweig, 1971).

Together, these insights emphasized the importance of energetic boundaries, but they left unresolved the broader problem of what ultimately limits the emergence of new trophic levels and why do they exist at all. Early work highlighted evolutionary and dynamical constraints on chain length (Hastings and Conrad, 1979, Pimm and Lawton, 1977), while comparative studies examined how productivity, disturbance, and ecosystem size might predict trophic length across ecosystems (Cohen et al., 2012, Pimm and Kitching, 1987, Post, 2002). Yet these approaches have struggled to identify general energetic or thermodynamic principles that set an upper bound (Borrelli and Ginzburg, 2014, Sterner et al., 1997). Three specific gaps stand out.

First, most work has emphasized *lower bounds* on supply (consumer thresholds), while the *possible upper bounds* compatible with producer persistence have not been clearly defined. Second, while dynamical models capture instability at large energy inputs, they do not translate into simple thermodynamic inequalities that bound trophic emergence (Hastings and Conrad, 1979, Rosenzweig, 1971). Third, environmental or physiological ceilings on energy input are often imposed *ad hoc*, without integration into a general framework (Brown et al., 2004, Sibly et al., 2012). This raises several new questions: What are the full set of energetic boundaries that constrain the emergence of trophic systems? How do these boundaries interact to determine whether additional trophic levels can persist? And can these ecological constraints be grounded explicitly in the thermodynamic laws that govern energy flow and dissipation?

In this paper, we develop a thermodynamically grounded framework to address these questions. We begin by formulating energetic balances for trophic chains of arbitrary length and showing how they can be expressed as feasibility conditions. We then use this formalism to identify three distinct types of energetic boundaries and to analyze their overlap as the condition for trophic emergence. Finally, we illustrate the implications of this framework with worked examples and derive a general criterion for the maximum length of trophic chains. This approach integrates classical ecological theory with fundamental thermodynamic principles, providing a new lens on the energetic limits of trophic systems.

### Energetic flow model and its feasibility

We represent ecosystems as populations linked by energy flows. For clarity, we focus here on strictly vertical trophic chains with *L* levels, indexed from the basal level (*i* = 1) to the top consumer (*i* = *L*). This setting provides a transparent baseline to illustrate the energetic constraints at play. However, the same logic applies to more complex interaction structures, including nonlinear interactions, food webs with branching, omnivory, or cross-feeding, as we show in the Appendix. The generality of the framework arises because the constraints we derive are a direct consequence of energy conservation and dissipation, and thus do not depend on a particular structure of population interactions (see Figure 1).

**Figure 1.**
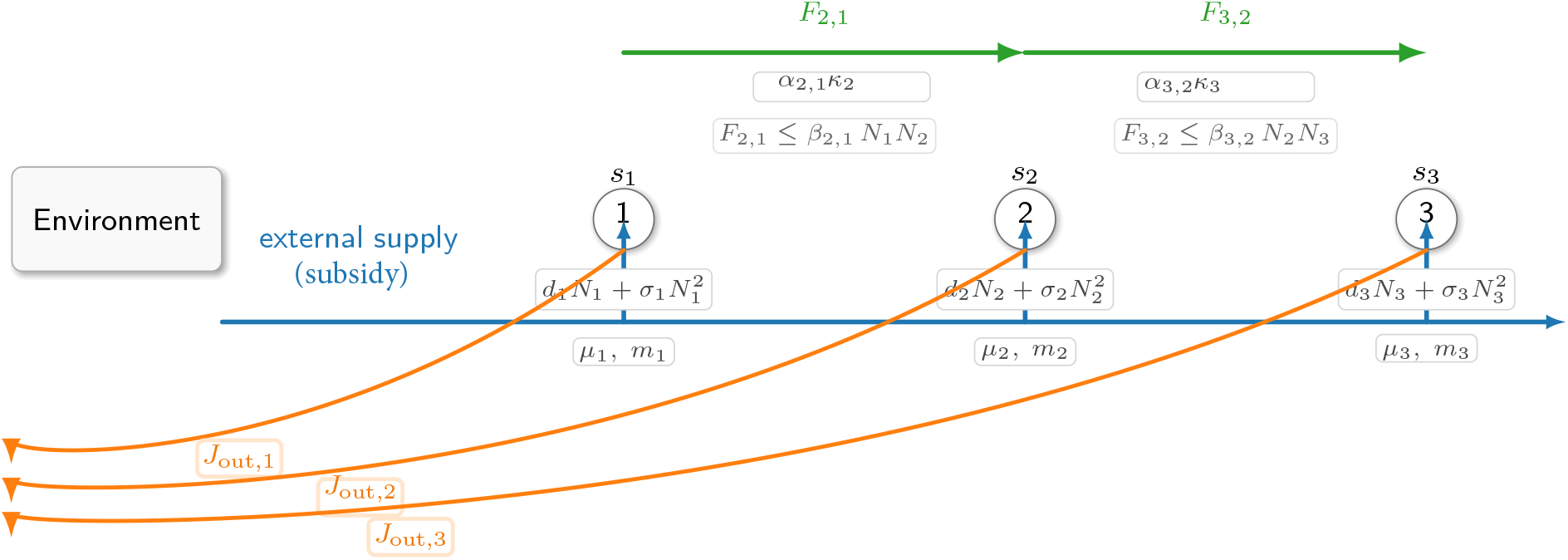
Ecosystem energy-flow schematic. A single blue flux represents the external energy reservoir feeding levels via taps *s*_*i*_ (arrowheads). Green arrows are trophic fluxes *F*_*i*+1,*i*_ from level *i* to *i*+1 along a dedicated track, annotated by the net conversion efficiency (*η*_*i*+1,*i*_ = *α*_*i*+1,*i*_*κ*_*i*+1_) (where *α*_*i*+1,*i*_ is the assimilation efficiency and *κ*_*i*+1_ is the trophic transfer efficiency) and by capacity constraints *F*_*i*+1,*i*_ ≤ *β*_*i*+1,*i*_*N*_*i*_*N*_*i*+1_. Orange curves are dissipative exports *J*_out,*i*_ returning to an environment. Circles mark trophic levels with biomass *N*_*i*_; per-level losses are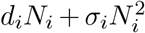; and *µ*_*i*_ (energy per biomass) with *m*_*i*_ (scaling) link biomass to energetic rates. All arrows indicate the positive direction of flow (flux = rate of energy transfer). At equilibrium, the biomass (energy) balance of trophic *i* requires that the total energy it assimilates from external inputs and prey consumption is at least as large as its internal energy demands and exports to consumers: 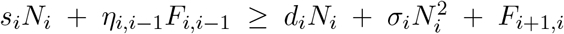.

To keep all terms dimensionally consistent and ecologically relevant, we work in biomass/time units, however, energy rates can be recovered by multiplying biomass by the specific energy content *µ*_*i*_ (kcal kg^−1^). Let *N*_*i*_ (kg) denote the biomass of population *i*. We assume that each individual in population *i* is characterized by a typical body mass *m*_*i*_ (kg), a per-biomass maintenance demand *d*_*i*_ (day^−1^), and a self-limitation parameter *σ*_*i*_ (day^−1^· kg^−1^) that captures energetic losses due to crowding. Populations may also assimilate energy from an external source (e.g., the Sun or other inorganic free energy source) at a per-biomass rate *s*_*i*_ (day^−1^), subject to an upper bound 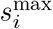imposed by environmental or physiological limits. We call this external flux *subsidy* (Polis et al., 1997). If *s*_*i*_ = 0, we call population *i* unsubsidized.

For consumers (*i* ≥ 2), a nonzero external term *s*_*i*_ *>* 0 can represent cross-feeding or detrital inputs to that level. Throughout, assimilation fractions *α*_*i*+1,*i*_ ∈ (0, 1) and *trophic transfer efficiencies κ*_*i*+1_ ∈ (0, 1) (fraction of assimilated prey that becomes consumer biomass) appear only via the product *η* = *α*_*i*+1,*i*_*κ*_*i*+1_ in feasibility bounds; we make this explicit here to reduce parameterization.

Energy can also flow between populations: a consumer at level *i*+1 extracts energy from level *i* at a rate *F*_*i*+1,*i*_ (kg day^−1^). Only a fraction *α*_*i*+1,*i*_ of this flux is assimilated, with the rest dissipated as heat, consistent with the Second Law of thermodynamics (Kondepudi and Prigogine, 2014, Prigogine, 1962). The flux is limited by an encounter/uptake rate coefficient *β*_*i*+1,*i*_ (day^−1^kg^−1^), leading to the capacity bound

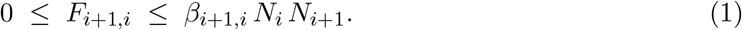

At equilibrium, the biomass (energy) balance (First Law of thermodynamics) of population *i* requires that the total energy it assimilates from external inputs and prey consumption is at least as large as its internal energy demands and exports to consumers:

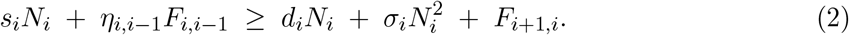

To determine whether a trophic chain can persist under a given subsidy vector **s** = (*s*_1_, …, *s*_*L*_), we require that every population maintains *at least a minimal threshold biomass*. For simplicity, we set this threshold to *N*_*i*_ ≥ *m*_*i*_, which means that each population should have at least one individual. This choice is arbitrary: any positive threshold would yield the same qualitative results, because the inequalities that define persistence simply rescale with the chosen values.

At the boundary *N*_*i*_ = *m*_*i*_ of this minimal threshold biomass, the condition of Eq. (2) gives

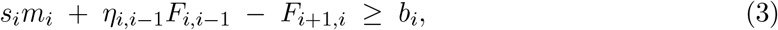

where *b*_*i*_ is the internal demand at the threshold *m*_*i*_:

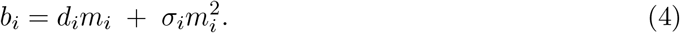

Equations (3-4) isolate the internal, flux-independent demand of level *i* (biomass/time) and let all boundary constraints take a uniform compact form, which we will use later in the text.

The set of all subsidy vectors **s** for which there exist nonnegative fluxes *F* (Eq. 1) satisfying Eq. (3) defines the set of environmental conditions under which the chain is energetically sustainable. We refer to this set as the *feasibility domain* (Logofet, 1993, Saavedra et al., 2017), denoted *𝒟*_*L*_. Formally, we write *D*_*L*_ for the set of supply vectors **s** for which *exactly L trophic levels* are feasible at equilibrium *N*^∗^(**s**):

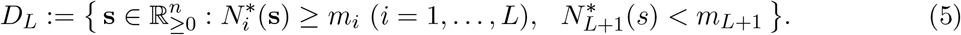

### Linking thermodynamic principles and feasibility constraints

The feasibility inequalities above specify when a chain can persist, but not which feasible subsidy conditions are most relevant. To address this, we introduce an optimization principle grounded in thermodynamics. First, consider the total external input (biomass throughput) at equilibrium

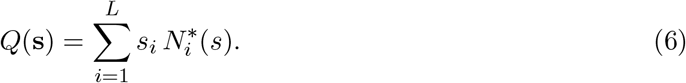

To connect with thermodynamics, we now switch to *energy* units by multiplying each biomass quantity by its specific energy content *µ*_*i*_ (kcal kg^−1^). To avoid confusion with the biomass throughput *Q*(**s**), we temporarily add a subscript *E* for energy. Define *G*_*E*_(*t*) (stored Gibbs free energy, kcal), *Q*_*E*_(*t*) (external input power, kcal day^−1^), and *J*_out,*E*_(*t*) (export to environment, kcal day^−1^) as follows:

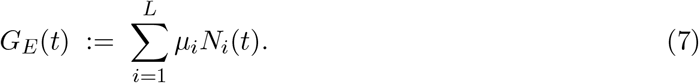

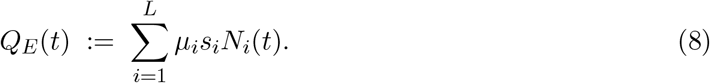

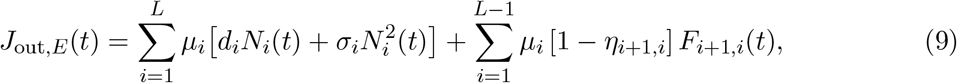

where only a fraction (0 *< η <* 1) of the assimilated energy (biomass) coming from trophic level *i* − 1 gets transformed into energy (biomass) of trophic level *i*.

Along solutions of the ecological dynamics and following the First Law of Thermodynamics,

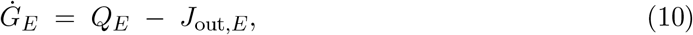

i.e., the rate of change of stored energy equals input power minus exported (power-dissipated) energy. Then, let *S*(*t*) be the system entropy and 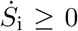 the irreversible entropy production. For an open isothermal system at environmental temperature *T*_*e*_ the Second Law gives

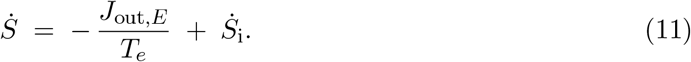

Substituting 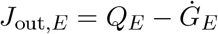 from (10) gives

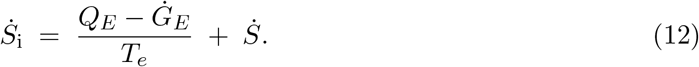

For any bounded *f* (*t*), define 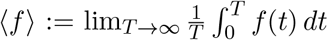 . Under permanence (i.e., each *N*_*i*_(*t*) is ultimately bounded above and below), *G*_*E*_(*t*) and *S*(*t*) are bounded, hence 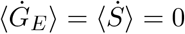 .

Averaging (12) yields

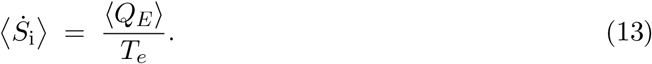

If the trajectory converges to an equilibrium *N*^∗^(**s**) (the assumption taken in the main text), then 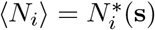 and

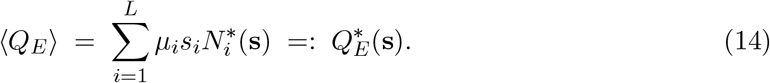

Therefore,

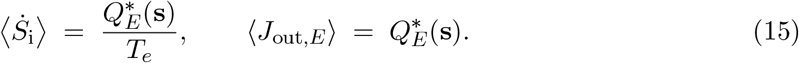

Since *µ*_*i*_ *>* 0 are fixed coefficients, maximizing 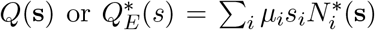 are equivalent optimization problems (up to a positive, level-weighted scaling). Hence maximizing *Q*(**s**) is thermodynamically justified by Eq. (15).

The optimization problem can therefore be written as

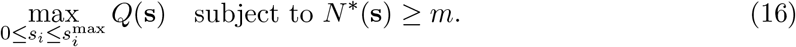

These conditions are sufficient to interpret *Q*(**s**) as the time-averaged external power input that is ultimately dissipated. They are not necessary laws for all ecosystems; when violated (e.g., strong matter export or non-isothermal gradients), the optimization should be read as a modeling assumption rather than a theorem.

This formulation provides the formal link between the *feasibility conditions* (Eqs. 3–4) and the *thermodynamic principle* of maximizing energy flow. Following this link, it is expected that the marginal value of basal input (i.e., biomass response to a unit increase of *s*_*i*_), ∂*Q/*∂*s*_1_, is always greater than or equal to that of any consumer input, ∂*Q/*∂*s*_*i*_ for *i* ≥ 2. Thus, the solution to the maximum output power is always at the unsubsidized axis

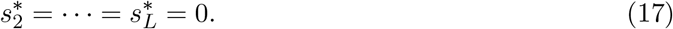

Ecologically, this expresses the principle of *subsidy inefficiency*: external energy to consumers directly never increases per-capita efficiency. This solution always allocates subsidy at the basal level alone. Direct subsidies to non-basal populations (*s*_*i*_ *>* 0 for *i* ≥ 2) are energetically inefficient: while they increase total input, each trophic transfer incurs a conversion loss (*κ <* 1), so system-wide output grows less per unit input than if the same energy were invested at the base. Trophic levels therefore emerge as the unique solution that maximizes energy throughput. Full mathematical details of this result are given in the Appendix.

### Energetic boundaries

The link between feasibility conditions and thermodynamic principles also makes explicit three energetic boundaries operating on the basal level (*i* = 1), which we denote by 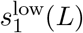 (lower bound), 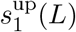 (internal upper bound), and 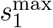 (external caps), as follows:

#### (i) Lower bound 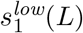 (consumer thresholds)

Consumers (*i* ≥ 2) must assimilate enough inflow to cover 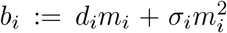. On the unsubsidized slice (*s*_*j*≥2_ = 0) and at feasibility (*N*_*i*_ = *m*_*i*_), define the cumulative minimal requirement *T*_*i*_ (kg day^−1^) as the least inflow that must arrive to level *i* from level *i* − 1 so that levels *i, i*+1, …, *L* can all be held at threshold. Set *T*_*L*+1_ = 0 and use the backward recursion

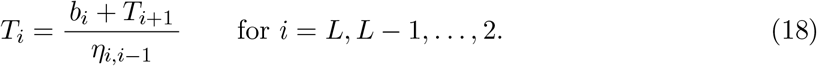

Holding the producer at *N*_1_ = *m*_1_, the most it can spare is 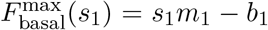. Feasibility on the unsubsidized slice is therefore equivalent to 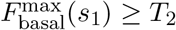, i.e.,

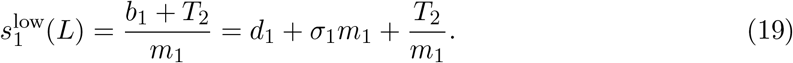

Unwinding the recursion yields the closed form

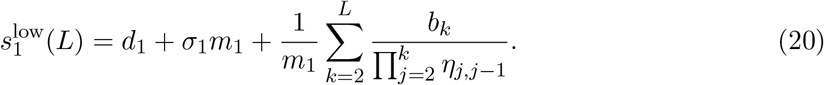

#### (ii) Basal upper bound 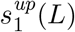 (protection limit)

At the producer boundary *N*_1_ = *m*_1_, the basal balance is

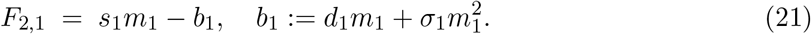

However, prey removal is encounter-limited:

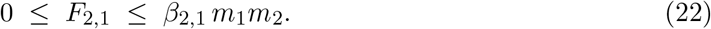

Feasibility therefore requires 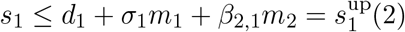. More generally, for length *L*,

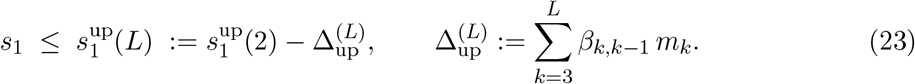

If 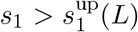, the producer would need a removal flux exceeding any biophysically attainable *F*_2,1_, and no state with *N* ≥ *m* can satisfy the boundary balances (First Law). This protection limit is an internal, encounter-driven cap on enrichment (a thermodynamic-style restriction on feasible throughput).

#### (iii) External/physiological caps 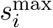

Environmental or physiological limits prevent supplies from increasing indefinitely. We encode these as box constraints on each coordinate:

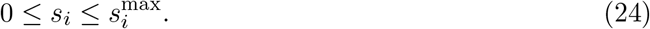

The intersection of these three constraints defines the bounded subset of supply space in which all populations can be maintained. We call this the *emergence interval* for a chain of length *L*:

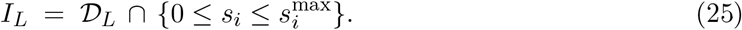

Equivalently, along the producer axis (no consumer subsidy), the interval is

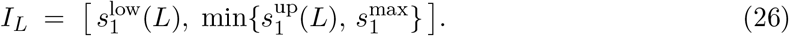

A new trophic level *L*+1 can emerge without direct subsidy precisely when the extended system remains feasible at *s*_*L*+1_ = 0, i.e.

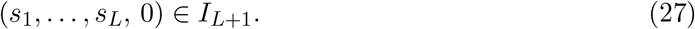

### Two trophic levels

To better illustrate our framework, we begin with the simplest trophic system of a producer (basal population 1) and a single consumer (population 2). At the persistence threshold, the feasibility inequalities can be written in linearized balance form

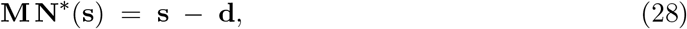

where 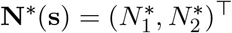 is the equilibrium biomass vector. For a two-level chain the balance matrix is tri-diagonal,

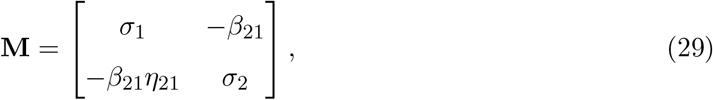

with positive diagonal entries (self-limitation) and negative off-diagonals (trophic couplings). The sign convention places the system in standard *M* -matrix form (Logofet, 1993). It is important to stress that this is not the direct algebraic form of the original dynamics, but a linearized balance representation constructed at the feasibility threshold. The purpose of this reformulation is to encode the feasibility inequalities in a way that makes the matrix **M** an *M* -matrix with positive diagonal entries and nonpositive off-diagonals. In this form, classical *M* -matrix theory ensures that **M**^−1^ ≥ 0, which guarantees monotone biomass responses to external supply (subsidy). Thus the transformation is a matter of analytical sign convention, not a modification of the underlying ecological dynamics (see Appendix for the derivation and proof of equivalence).

Feasibility requires **M** to be an *M* -matrix with nonnegative inverse (see Appendix), which here reduces to

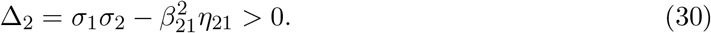

We maximize total input 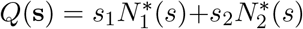 over the feasible set 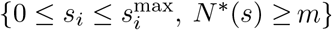. As we noted before, by the subsidy-inefficiency result (see Appendix), the maximizer lies on the *unsubsidized axis* with 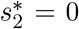 (and if 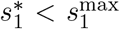 then necessarily 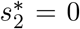). On the unsubsidized axis **s** = (*s*_1_, 0), feasibility at the thresholds *N*_1_ = *m*_1_, *N*_2_ = *m*_2_ requires that the producer can meet its own needs and supply enough energy to the consumer. This translates into several energetic bounds.

#### Producer-alone requirement

With no consumer extraction, the producer balance 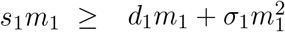 yields the basal requirement

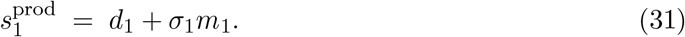

#### Consumer threshold

At the consumer threshold (*N*_2_ = *m*_2_) and *s*_2_ = 0, the consumer requires 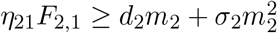. The minimal required inflow is

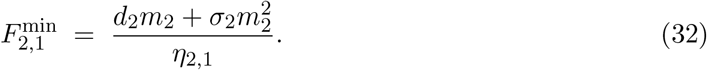

From the producer balance 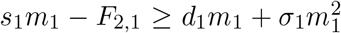, the maximum flux the producer can supply at given *s*_1_ is 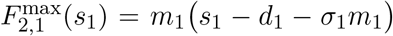. Requiring 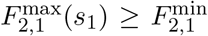 gives the minimal basal supply that allows the consumer to persist:

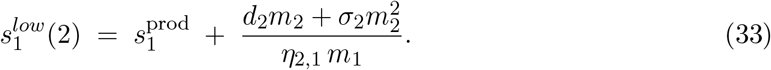

Ecologically, Eq. (33) shows that the minimal basal input required for the consumer combines two components: (i) the producer’s own requirement to cover maintenance and self-limitation, *d*_1_ + *σ*_1_*m*_1_, and (ii) the additional input needed to support the consumer at its threshold, scaled by the assimilation efficiency *η*_2,1_ and distributed over the producer biomass *m*_1_.

#### Basal protection limit

Even if the producer can in principle allocate large flux, encounters cap the trophic flow at *F*_2,1_ ≤ *β*_2,1_*m*_1_*m*_2_. Imposing 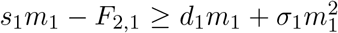 at this cap yields an *upper* feasible basal input:

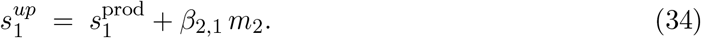

Ecologically, the above equation is *analogous in spirit* to the classic paradox of enrichment: at very high basal input, consumers can extract more energy than producers can maintain, driving producers below their threshold.

The emergence interval for two levels on the unsubsidized axis is therefore

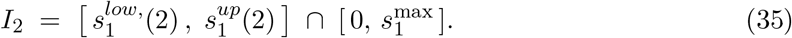

#### Feasibility vs. emergence

A nonempty feasible region in (*s*_1_, *s*_2_) does not by itself guarantee a nonempty unsubsidized emergence interval *I*_2_; the latter requires the basal-axis slice to intersect the feasible set.

The range is nonempty only if the maximal encounter capacity of the consumer is sufficient to cover its energetic demand, that is,

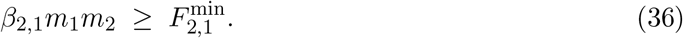

If this inequality fails, then even under arbitrarily large basal input the consumer cannot obtain enough energy, because the rate of encounters places a hard ceiling on trophic flux. Ecologically, this condition expresses the requirement that consumer foraging capacity must be high enough relative to its metabolic and crowding costs; otherwise the trophic link cannot be established, and the chain collapses to a single level. Figure 2 presents a worked example.

**Figure 2.**
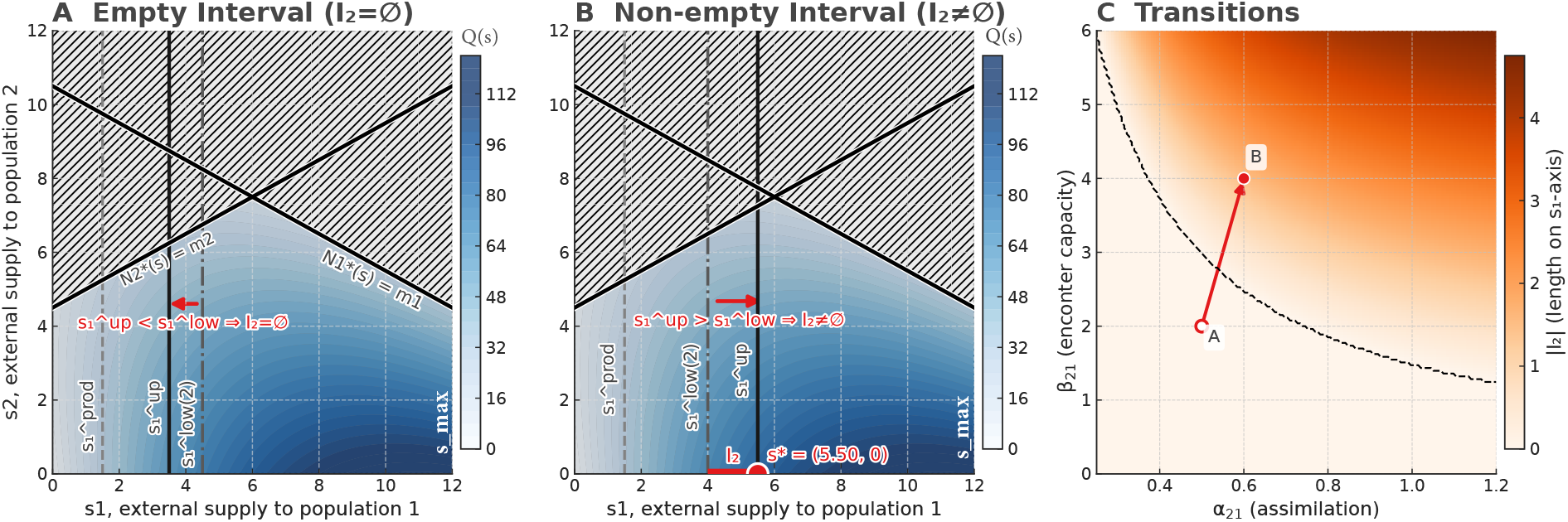
Feasibility, emergence interval, and optimality in a two-level chain. Panels (A-B) show the external supply, subsidy, (*s*_1_, *s*_2_) plane with feasibility boundaries 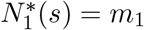 and 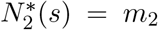 (black), the infeasible region (hatched), the external cap *s*_*max*_ We mask values of *Q*(**s**) outside the feasible set (values *<* 0 are suppressed) so the color scale reflects feasible throughput only, and a heat map of biomass-throughput cap 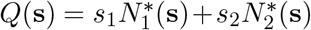. We use the shared parameters *m*_1_ = *m*_2_ = 1, *d*_1_ = *d*_2_ = 1, *σ*_1_ = *σ*_2_ = 0.5, and 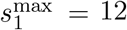 . Axis bounds on the *s*_1_-axis are 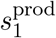 (producer-alone requirement), 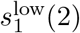 (consumer threshold), 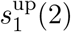 (basal protection limit). The emergence interval (for a two-level chain) is 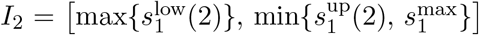 With *η*_21_ = 0.50 and *β*_21_ = 2 we obtain 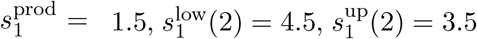, so *I*_2_ = ∅ (no emergence). No optimizer is shown. That is, while the panel shows a nonempty feasible region in (*s*_1_, *s*_2_) that happens when feasibility requires *s*_*i*_ *>* 0, it has an infeasible region (empty emergence interval, *I*_2_ = ∅) at the unsubsidized slice (*s*_2_ = 0) that does not intersect the feasible set. This is not a contradiction; it distinguishes overall feasibility from unsubsidized emergence. **(B)** With *η*_21_ = 0.60 and *β*_21_ = 4.00 we obtain 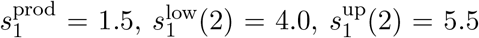, hence *I*_2_ = [4.0, 5.5] (non-empty). The maximizer of *Q*(**s**) lies on the unsubsidized axis 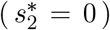 at 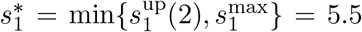 (red point and segment *I*_2_), illustrating “subsidy inefficiency.” Note that we draw 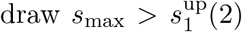, indicating thermodynamic constraint. However, this can also be drawn 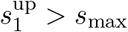, indicating physiological or environmental constraint. **(C)** Parameter plane (*η*_21_, *β*_21_) colored by the emergence-interval length |*I*_2_| (white = 0, darker = larger |*I*_2_|). Markers show the (A) and (B) settings and the transition that opens the emergence interval *I*_2_.

### Three trophic levels

By adding a third trophic level 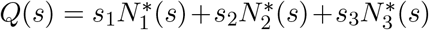 leads to the same feasibility and optimization criterion and subsidy inefficiency argument 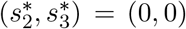: the optimizer concentrates all subsidy at the base (see Appendix). Accordingly, we analyze thresholds on the basal axis (*s*_1_, 0, 0), from which the new emergence interval *I*_3_ is read. Adding a secondary consumer (*i* = 3) creates two consumer thresholds and a tighter basal protection limit.

*Consumer thresholds*. Level 2 requires

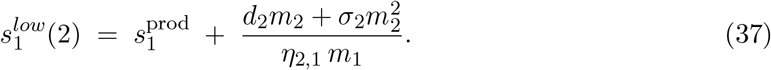

Level 3 requires 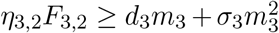, i.e., a minimal flux 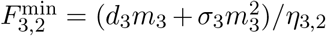, which must ultimately be sourced from producer input. Projecting this need back to the base gives

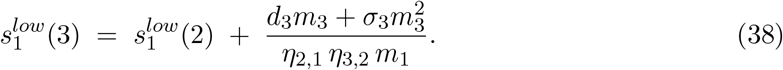

Ecologically, this stacks the costs of levels 2 and 3, discounted by the successive assimilation efficiencies *η*_2,1_*η*_3,2_ and apportioned to the basal biomass *m*_1_.

#### Basal protection limit

At three levels, maximal extraction from level 1 is *F*_2,1_ ≤ *β*_2,1_*m*_1_*m*_2_, but the presence of level 3 increases total exports through level 2. This tightens the basal protection limit to

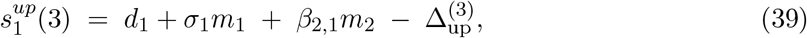

where 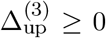 represents the additional effective export induced by the secondary consumer (Appendix). Thus, adding a higher consumer both raises the lower thresholds and reduces the upper limit.

The emergence interval then becomes

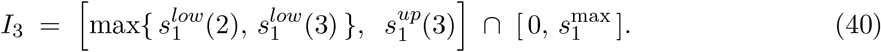

For three levels, *I*_3_ is nonempty only if both encounter constraints are satisfied:

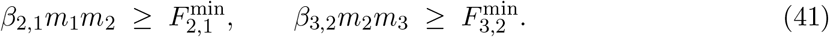

Violation of either inequality implies that the corresponding consumer cannot obtain enough energy even under arbitrarily high basal input *s*_1_. Ecologically, this means that each trophic link must provide sufficient throughput relative to the consumer’s energetic costs; if foraging efficiency is too low at any step, the chain collapses to a shorter length. Figure 3A presents a worked example.

**Figure 3.**
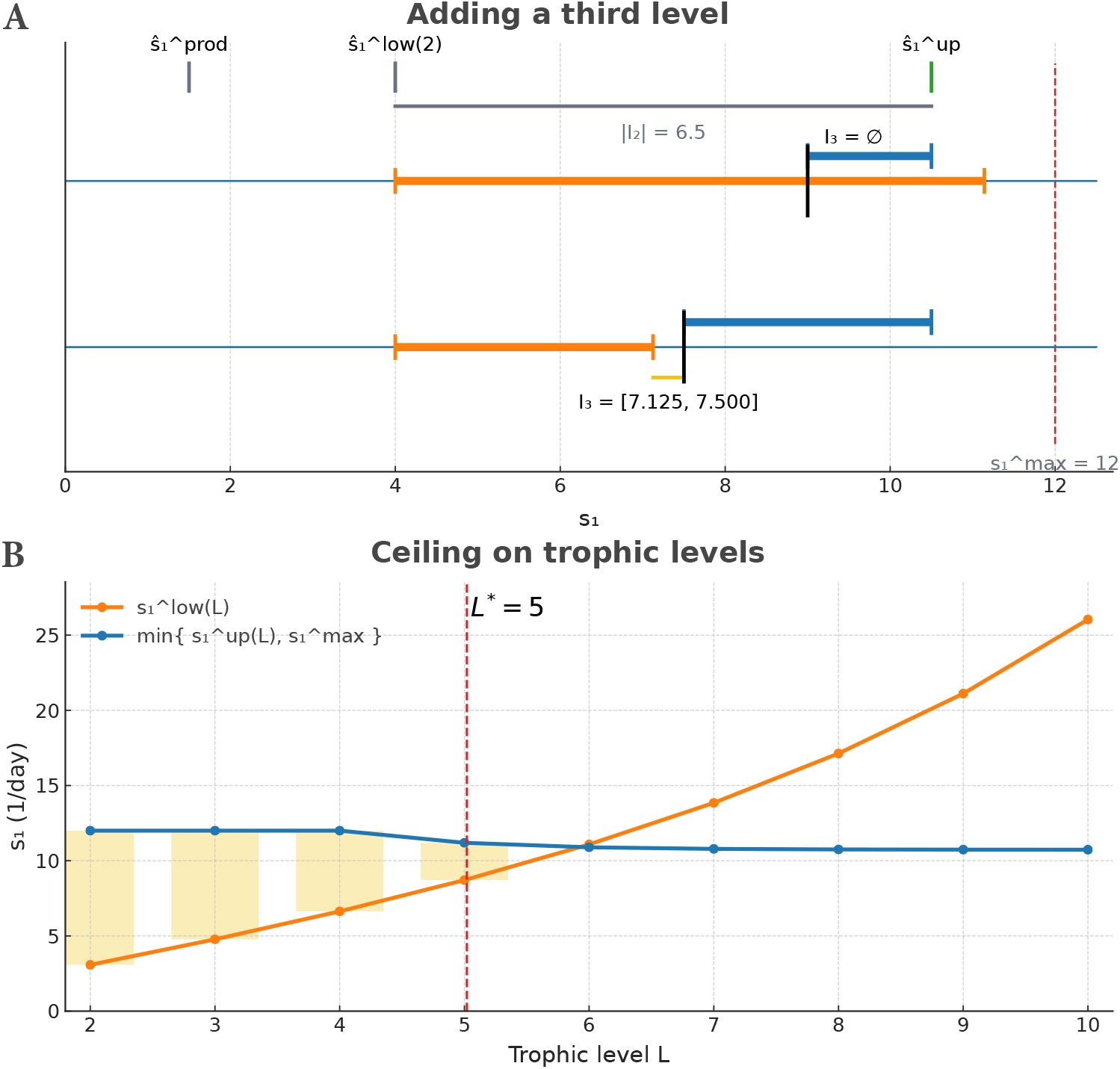
Adding and limiting trophic levels. We fix the external cap 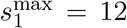and, unless noted, set *m*_*i*_ = *d*_*i*_ = 1 and *σ*_*i*_ = 0.5. Along the basal axis the three bounds are the stacked-consumer lower bound 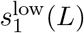, the basal-protection upper bound 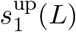, and the external cap 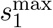. The unsubsidized emergence interval is 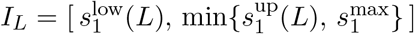, with 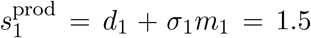 . *Baseline (two levels):* with *η*_2,1_ = 0.60 and *β*_2,1_ = 9.00 we obtain 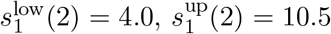, so |*I*_2_| = 6.5. **A**: Adding a third level. For the link 3 → 2, 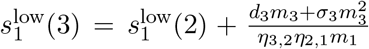 and 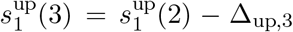 with Δ _up,3_= *β*_3,2_ *m*_3_ . Top case (*η*_3,2_ = 0.35, *β*_3,2_ = 1.50): Δ_low,3_ = 1.5*/*(0.35 0.60) 7.143, Δ_up,3_ = 1.50, hence *I*_3_ = ∅. Bottom case (*η*_3,2_ = 0.80, *β*_3,2_ = 3.00): Δ_low,3_ = 3.125, Δ_up,3_ = 3.0, hence *I*_3_ = [7.125, 7.5]. In both cases the optimizer lies on the unsubsidized axis with 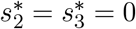; black ticks mark 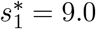 (top) and 7.5 (bottom). Colors: 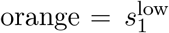 and lower-bound shifts; blue = 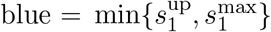 and upper-bound shifts; yellow = feasible intervals. **B**: Ceiling with increasing trophic levels. For *L* = 2, …, 10 we plot 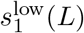 and 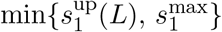and shade |*I*_*L*_| (yellow). To illustrate attenuation with height we use 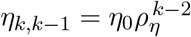 and 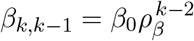 with *η*_0_ = 0.95, *β*_0_ = 20.00, *ρ*_*η*_ = 0.98, *ρ*_*β*_ = 0.35. For these values |*I*_*L*_| *>* 0 for *L* ≤ 5 and is empty at *L* = 6, i.e., *L*^∗^ = 5.

### Ceiling on trophic levels

Generally, for a chain of arbitrary length, each consumer imposes a lower bound on basal input, while the basal producer population imposes a possible upper bound, and the environment sets subsidy caps. The emergence interval is

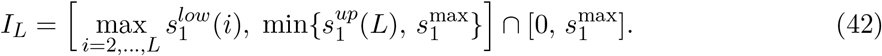

As *L* increases, consumer thresholds 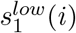 rise due to compounding maintenance demands, while 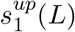 falls as basal producers become increasingly vulnerable to cumulative exports. This opposing movement causes the emergence interval to contract with chain length. The maximum sustainable trophic length is therefore

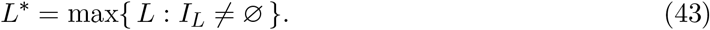

## Discussion

Food webs have long been recognized as the backbone of ecosystem organization, yet the question of what limits their vertical extent—how many trophic levels can coexist—has remained one of the most persistent puzzles in ecology (Pimm, 1982). For more than half a century, ecologists have debated whether food-chain length is controlled primarily by productivity, disturbance, ecosystem size, or evolutionary stability (Cohen et al., 2012, Hastings and Conrad, 1979, Pimm and Kitching, 1987, Pimm and Lawton, 1977, Post, 2002). These studies established a rich set of insights: resource competition theory provided the concept of a minimum requirement (*R*^∗^) for consumer persistence (Tilman, 1980), structural and matrix-based approaches formalized feasibility domains as a way to capture lower energetic bounds (Logofet, 1993, Loreau, 2010, Saavedra et al., 2017), and consumer–resource models revealed the paradox of enrichment, where excess supply destabilizes producers (Rosenzweig, 1971). Still, none of these approaches have yielded a general thermodynamic principle specifying why food chains are almost always short, or what determines the absolute ceiling to trophic length (Borrelli and Ginzburg, 2014, Sterner et al., 1997).

Our results show that this long-standing problem can be reformulated in terms of a geometry of feasibility constrained by the laws of thermodynamics. By embedding energetic balances into a throughput-maximizing principle (Eq. 16), we identify a triad of energetic boundaries that jointly determine whether trophic levels can persist. Consumer thresholds provide lower bounds, ensuring each higher-level population receives enough upstream energy to maintain at least one individual—directly extending the *R*^∗^ logic of resource competition into food chains of arbitrary length (Logofet, 1993, Tilman, 1980). Basal protection limits impose possible upper bounds, because increasing basal input raises consumer extraction and dissipation until producers themselves are no longer viable. This reframes the paradox of enrichment (Rosenzweig, 1971) as a thermodynamic constraint. Finally, external supply caps represent environmental or physiological ceilings (Brown et al., 2004, Sibly et al., 2012), ensuring that total input remains finite.

The intersection of these three boundaries defines the emergence interval within which a chain of length *L* can exist. This perspective highlights how trophic persistence is not governed by a single limiting factor, but by the geometry of interacting constraints. Productivity or resource availability can shift the consumer thresholds downward, making higher levels more accessible, but may also tighten basal protection limits, reducing producer viability. Ecosystem size or environmental forcing determines the magnitude of the external cap, which can truncate the feasible range even when internal energetics would allow longer chains. In this sense, comparative observations of food-chain length across productivity or disturbance gradients (Pimm and Kitching, 1987, Post, 2002) can be reinterpreted as reflections of how ecological and thermodynamic boundaries overlap.

A striking consequence of our framework is the principle of subsidy inefficiency. The optimization analysis (Eq. 16) shows that external input is never directed toward non-basal populations (*s*_*i*_ = 0 for *i* ≥ 2). This result follows from the fact that the marginal system-wide return of basal input always exceeds that of non-basal subsidies, since every trophic transfer dissipates a fraction of energy. Ecologically, this means that supporting non-basal populations directly is never the most efficient way to sustain a trophic system. Instead, energy invested at the base uniquely maximizes system throughput, forcing the optimizer onto the unsubsidized axis. Trophic levels thus emerge not by assumption, but as the only energetically consistent configuration that reconciles feasibility with thermodynamic laws.

Generalizing to chains of arbitrary length, we find that the emergence interval contracts as trophic levels accumulate. Each additional consumer imposes a new lower threshold, while the basal protection limit falls under increasing cumulative exports. When these two internal constraints cross, the emergence interval vanishes: no basal input can simultaneously satisfy producers and consumers. This crossing defines a thermodynamic ceiling on food-chain length, a boundary beyond which further trophic extension is impossible. Crucially, while ecological limitations (such as external caps) can be relaxed by increasing supply, the thermodynamic ceiling is absolute: dissipative losses compound inexorably with trophic depth, and no amount of external input can remove this bound. This result provides a fundamental energetic explanation for why ecosystems never sustain indefinitely long chains.

Although we illustrated our approach with linear trophic chains for clarity, the underlying logic extends to more complex food-web architectures, including nonlinear dynamics, branching, and omnivory (Appendix). Because the constraints arise directly from energy conservation and dissipation, the triad of boundaries and the principle of subsidy inefficiency are robust features of energetically consistent systems. Moreover, the qualitative results do not depend on a particular optimization criterion: any objective consistent with the Second Law leads to the same axis-dominated solutions and the same progressive narrowing of the emergence interval.

Our framework highlights a direct link between ecological feasibility and nonequilibrium thermodynamics. The total external input, which we maximize, is mathematically equivalent to long-term entropy production (Kondepudi and Prigogine, 2014, Odum, 1988, Prigogine et al., 1973, Solé et al., 2024). Earlier formulations of the Maximum Entropy Production Principle (MEPP) proposed that ecosystems maximize entropy production within feasible states (Ulanowicz, 1987). Recent work has developed the Minimum Environmental Perturbation Principle (MEPP), showing that equilibria in consumer–resource models can also be understood as minimizing deviations from the unperturbed environment (Marsland et al., 2020). Our results complement these perspectives: whereas the MEPP addresses equilibrium selection *within* the feasible region, our analysis shows that thermodynamic principles also act on the *boundaries of feasibility*, dictating the very conditions under which trophic levels can emerge. This synthesis suggests a new way of understanding the organization of ecosystems: while ecological processes shape the position of energetic thresholds, thermodynamic laws ultimately impose an inescapable ceiling on trophic length. There is no escape from this ceiling—ecosystems may shift their boundaries under changing environments, but they cannot increase trophic levels indefinitely in defiance of the fundamental constraints imposed by dissipation.

## Acknowlegments

SS acknowledges support from the National Science Foundation under Grant No. DEB-2436069 and MIT Google Program for Computing Innovation. PM acknowledges support from grant ANID-Exploración 13220168, FB210005, BASAL funds for centers of excellence from ANID-Chile, and the ICTP through the Associates Programme and the Simons Foundation through grant number 284558FY19. SAL acknowledges support from the National Science Foundation under grant No. DMS-1951358 and ARO grant W911NF2410126. RPR acknowledges support from the Swiss National Science Foundation under CRSII5_202290, and 320030L-227556. MTA acknowledges support from UNAM-PAPIIT IA102225.

## Appendix

### M-matrix: unit-consistent linearization, sign standardization, and invariance

#### Model ingredients and symbols

For a trophic chain with levels *i* = 1, …, *L* (producer 1 to top consumer *L*):

- *N*_*i*_ : biomass of population *i* (kg).
- *m*_*i*_ *>* 0 : *threshold* biomass (kg) used to test feasibility (*N*_*i*_ ≥ *m*_*i*_).
- *s*_*i*_ : external supply (per-biomass rate, day^−1^).
- *d*_*i*_ : maintenance (per-biomass rate, day^−1^).
- *σ*_*i*_ *>* 0 : self-limitation (crowding) coefficient (kg^−1^ day^−1^) so that 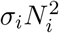 is a loss (kg day^−1^).
- *β*_*i*+1,*i*_ ≥ 0 : encounter/consumption coefficient (kg^−1^ day^−1^).
- *α*_*i*+1,*i*_ ∈ (0, 1) : assimilation fraction.
- *κ*_*i*+1_ ∈ (0, 1) : trophic transfer efficiency (dimensionless), mapping assimilated prey biomass into consumer biomass in the linearization.
- *η*_*i*+1,*i*_ ∈ (0, 1) : net conversion efficiency, defined by *η*_*i*+1,*i*_ := *α*_*i*+1,*i*_*κ*_*i*+1_. We use *η* in feasibility and threshold formulas.
- *µ*_*i*_ *>* 0 : energy content (kcal kg^−1^)

#### Energetic ODEs (two levels)

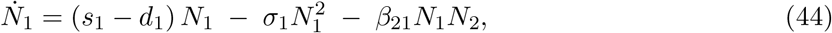

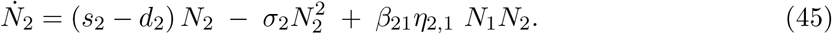

Here the producer loses to consumption, while the consumer gains from prey.

#### Boundary linearization at the feasibility threshold

We test feasibility by requiring *N*_*i*_ ≥ *m*_*i*_. On the boundary *N*_*i*_ = *m*_*i*_. We linearize all quadratic/bilinear terms by freezing *one* factor at its threshold:

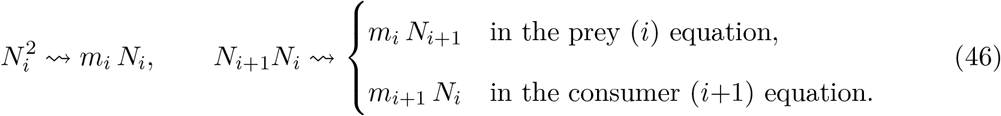

We also evaluate the maintenance term at the threshold: (*s*_*i*_ − *d*_*i*_)*N*_*i*_ ⇝ *s*_*i*_*N*_*i*_ − *d*_*i*_*m*_*i*_. This removes the multiplicative *N*_*i*_ from the RHS, yielding a linear balance in (*N*_1_, …, *N*_*L*_) with a *constant* RHS.

For two levels this gives

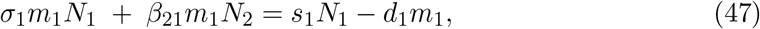

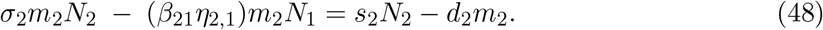

Dividing (47) by *m*_1_ and (48) by *m*_2_, and (at the boundary) replacing the remaining *s*_*i*_*N*_*i*_*/m*_*i*_ by *s*_*i*_, yields the per-biomass, unit-consistent form

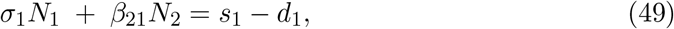

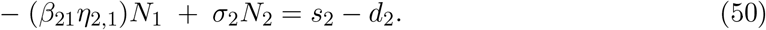

#### Sign standardization (M-matrix form)

We adopt the conventional *M* -matrix sign pattern by moving all inter-level terms to the left so that diagonal entries are positive and off-diagonals are nonpositive. Define the positive coefficients

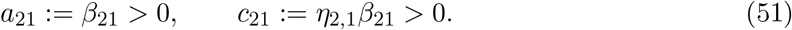

Then (49)–(50) can be sign-standardized as

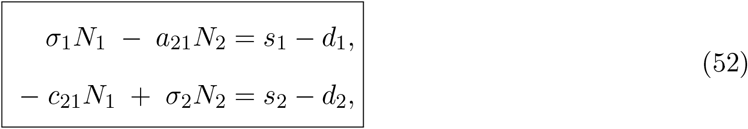

or, compactly, **MN** = **s** − **d** with

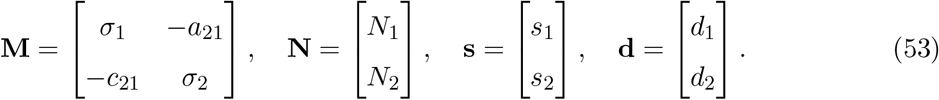

Every term in **MN** = **s** − **d** is in day^−1^. Entries of **M** are kg^−1^day^−1^, so **MN** is day^−1^.

#### General L and tri-diagonal structure

Repeating the same steps for *L >* 2 yields

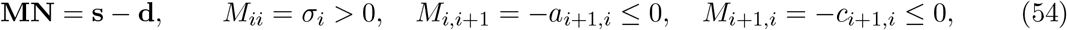

where

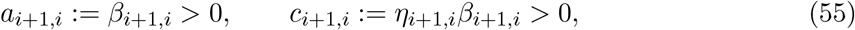

and **M** is tri-diagonal because only adjacent levels interact.

With *σ*_*i*_ *>* 0 and nonpositive off-diagonals, **M** is a (strictly) diagonally dominant *Z*-matrix; under the simple feasibility condition that all leading principal minors are positive (explicitly given in the main text for *L* ≤ 3), **M** is a nonsingular *M* -matrix and **M**^−1^ ≥ 0 (see Berman and Plemmons, 1994). This ensures monotone biomass responses to supply and allows the geometric arguments (feasibility domain, emergence interval) to proceed.

#### Invariance of feasibility under sign standardization

Let 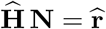 denote *any* boundary-linearized system obtained from (44)–(45) by freezing one factor at *m*_*i*_ in each quadratic/bilinear term and evaluating (*s*_*i*_ − *d*_*i*_)*N*_*i*_ at the threshold. Let **M** be the sign-standardized matrix defined above, obtained by moving all inter-level terms to the left. Then:

##### Lemma 1

(Feasibility invariance). *For any fixed m* = (*m*_1_, …, *m*_*L*_), *the following are equivalent:*

1. *There exists N* ≥ *m and nonnegative fluxes consistent with the linearization such that* 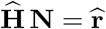.
2. *There exists N* ≥ *m such that* **MN** = **s** − **d**.

*Hence, the feasibility domain (set of* **s** *admitting* **N** ≥ **m***) and its induced emergence interval are unchanged by sign standardization*.

*Proof*. Both 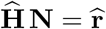 and **MN** = **s** − **d** are obtained from the same two operations: (i) boundary linearization (freezing one factor at *m*), and (ii) rearrangement of terms between sides of the equality. Operation (i) fixes coefficients; (ii) only changes signs of terms that are moved from the RHS to the LHS (or vice versa). In particular, there exists a diagonal matrix **S** with entries *±*1 (depending on whether a given inter-level term is kept on the RHS or moved to the LHS) such that 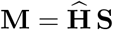 and 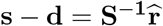. Because **S** is invertible and diagonal, the set of (**N, s**) pairs satisfying one system bijects to those satisfying the other with the *same* inequality **N** ≥ **m**. Therefore, the set of *s* for which a solution **N** ≥ **m** exists is identical in both representations.

*Notes on coefficients and units*. The numerical values of *a*_*i*+1,*i*_ and *c*_*i*+1,*i*_ depend on how constant factors are grouped when dividing by *m*_*i*_ and moving terms; their *signs* are invariant and all that is needed for the *M* -matrix structure. Using the per-biomass RHS (*s*_*i*_ − *d*_*i*_) keeps all terms in biomass / time and avoids mixing units such as (*s*_*i*_ − *d*_*i*_)*m*_*i*_. These properties justify the monotone dependence of **N**^∗^(**s**) on *s* and the geometric arguments used in the main text.

### Subsidy inefficiency

Recall **N**^∗^(**s**) = **As** + **h** with **A** = **M**^−1^ ≥ 0 and **h** = −**M**^−1^*d*, where **s** = (*s*_1_, …, *s*_*n*_)^*T*^ collects per-biomass subsidy rates and **d** = (*d*_1_, …, *d*_*n*_)^*T*^ collects per-biomass maintenance rates. Then

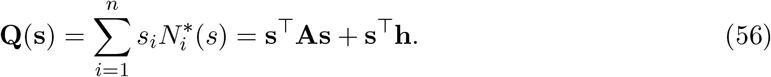

We work under the standard trophic-chain assumptions: *σ*_*i*_ *>* 0, *m*_*i*_, *µ*_*i*_ *>* 0, and *a*_*i*+1,*i*_ := *β*_*i*+1,*i*_, *c*_*i*+1,*i*_ := *η*_*i*+1,*i*_*β*_*i*+1,*i*_ (see previous Appendix Section).

#### Lemma 2

(Column–1 dominance for chain *M* -matrices). *Let* **v**^(*i*)^ := **Ae**_**i**_ *denote column i of* **A** = **M**^−1^, *i.e. the biomass response to a unit increase of s*_*i*_. *For the tri-diagonal trophic matrix* **M** *(sign-standardized as in the previous subsection)*,

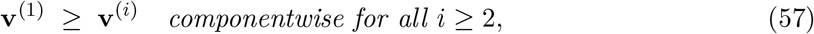

*with strict inequality in at least one component whenever any κ*_*j*_ *<* 1.

. Write the linear systems **Mv**^(*i*)^ = **e**_**i**_ explicitly:

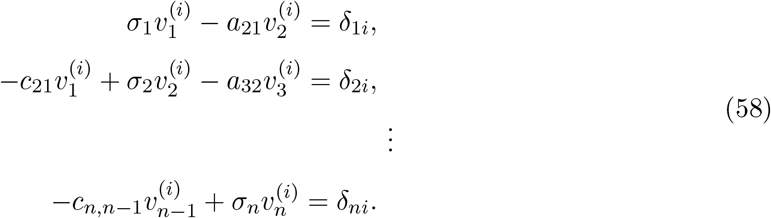

Solving the tridiagonal system **Mv** = **e**_**i**_ by forward/backward recursions yields continued-fraction expressions whose coefficients are products of the dimensionless ratios *θ*_*k*+1,*k*_ := *a*_*k*+1,*k*_*/σ*_*k*+1_ and *ϕ*_*k,k*−1_ := *c*_*k,k*−1_*/σ*_*k*_. For each adjacent pair we have *θ*_*k*+1,*k*_ *ϕ*_*k*+1,*k*_ = (*a*_*k*+1,*k*_*c*_*k*+1,*k*_)*/*(*σ*_*k*+1_*σ*_*k*_) *<* 1, because the 2 *×* 2 principal minors satisfy *σ*_*k*_*σ*_*k*+1_ − *a*_*k*+1,*k*_*c*_*k*+1,*k*_ *>* 0. Hence, away from the forcing index *i*, column entries attenuate geometrically by factors *<* 1. Column 1 has the shortest attenuation path to every level; any column *i* ≥ 2 incurs at least one extra lossy step, giving **v**^(1)^ ≥ **v**^(*i*)^ componentwise, with strict inequality whenever some *a*_*k*+1,*k*_*c*_*k*+1,*k*_ *>* 0 (equivalently, some transfer efficiency is *>* 0). Here *a*_*k*+1,*k*_ = *β*_*k*+1,*k*_ and *c*_*k*+1,*k*_ = *α*_*k*+1,*k*_*κ*_*k*+1_*β*_*k*+1,*k*_ with 0 *< η*_*k*+1,*k*_ *<* 1.

For each supply component *s*_*i*_, the *marginal value* is the partial derivative

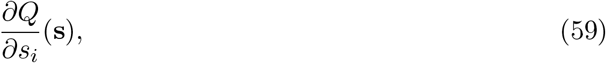

which measures the increase in system-wide output *Q*(**s**) per infinitesimal increase in *s*_*i*_. We call 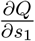 the *basal marginal value*, and 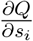 for *i* ≥ 2 the *consumer marginal values*.

#### Proposition 1

(Axis optimality under trophic M-matrices). *For Q*(**s**) = **s**^*T*^**As** + **s**^*T*^**h** *with* **A** = **M**^−1^ ≥ 0 *and* 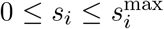, *there exists a maximizer on the unsubsidized axis, i.e*., *with s*_2_ = · · · = *s*_*n*_ = 0. *Moreover, if* 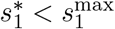 *at a maximizer, then necessarily* 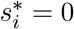 *for all i* ≥ 2.

#### Remark 1

(Scope of axis optimality). *Proposition 1 characterizes optimality* subject to feasibility. *When feasibility requires s*_*i*_ *>* 0 *for some consumers (e.g*., *due to caps or thresholds in Fig. 2A), the maximizer may lie on a subsidized boundary. The axis solution* 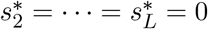 *holds on the unsubsidized feasible slice; under additional caps it remains the* direction *of improvement, but the optimizer can be pinned at a corner of the feasible polytope*.

*Proof*. Let *v*^(*i*)^ := **Ae**_**i**_ be column *i* of **A** = **M**^−1^. By Lemma 2 (column–1 dominance), **v**^(1)^ ≥ **v**^(*i*)^ componentwise for all *i* ≥ 2. Consider any feasible **s** with some *s*_*i*_ *>* 0 (*i* ≥ 2). For a small *ε >* 0, compare **s** to 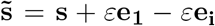 (staying within the box). Then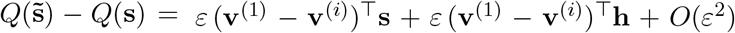. The first term is nonnegative because **s** ≥ 0 and **v**^(1)^ ≥ **v**^(*i*)^, while the second is nonpositive since **h** ≤ 0. If the net first-order change is nonnegative, *Q* does not decrease under this reallocation; if it is negative, we can move along coordinate directions to a vertex of the box and repeat the exchange argument. By compactness, an optimizer exists; choose one with the fewest positive consumer subsidies. The exchange argument and Lemma 2 then imply that all consumer subsidies vanish at an optimizer.

#### Corollary 1

(Axis optimality and vanishing consumer subsidies). *Let* **s**^⋆^ *be any maximizer of Q over the feasible box and the feasibility region* **N**^∗^(**s**) ≥ **m**. *Then*

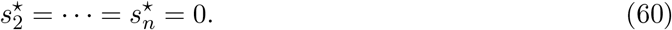

*Moreover, if* 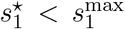, *the Karush-Kuhn-Tucker (KKT) conditions (Bertsekas, 1999) force strict equality and the maximizer lies on the* unsubsidized axis, *defined as*

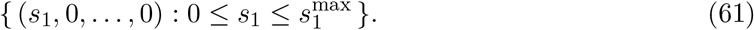

*Any vector with s*_*i*_ *>* 0 *for some i* ≥ 2 *lies on a* subsidized axis *and is dominated in Q by reallocating that subsidy to the basal input*.

Lemma 2 formalizes *subsidy inefficiency*: because each trophic transfer is lossy (*κ <* 1) and dissipation is positive (*σ >* 0), a unit of energy injected at a consumer produces a weaker system-wide response than the same unit injected at the base. Proposition 1 shows that the basal marginal value ∂*Q/*∂*s*_1_ always dominates any consumer marginal value ∂*Q/*∂*s*_*i*_ (*i* ≥ 2). The optimization therefore concentrates supply on the unsubsidized axis, and trophic emergence occurs only if this axis intersects the feasibility domain. If the axis lies entirely outside the feasible set, the optimizer is forced to a corner set by the external caps and no unsupplied trophic emergence is possible.

### Robustness and generality beyond linear chains

Recall **N**^∗^(**s**) = **As** + **h** with **A** = **M**^−1^ ≥ 0 and **h** = −**M**^−1^*d*, where **s** = (*s*_1_, …, *s*_*n*_)^*T*^ collects per-biomass subsidy rates and **d** = (*d*_1_, …, *d*_*n*_)^*T*^ collects per-biomass maintenance rates. So far we have assumed a strict chain structure (1 → 2 → · · · → *n*), in which each consumer feeds only on the immediately lower level. We now relax this assumption and allow consumers to feed on multiple lower levels (omnivory), yielding a directed acyclic trophic graph (DAG).

At the feasibility boundary, we linearize the bilinear feeding terms by freezing one factor at its threshold value. If consumer *i* has prey set 𝒫_*i*_ ⊂ {1, …, *i* − 1} and prey *k* has consumer set *C*_*k*_ := { *i* : *k* ∈ 𝒫_*i*_ }, the balance equations retain

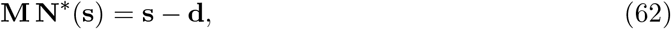

with diagonal entries *M*_*ii*_ = *σ*_*i*_, and off–diagonal entries

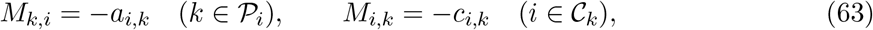

where

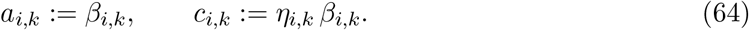

By construction, every *M*_*ij*_ has units kg^−1^ day^−1^, so **MN**^∗^(**s**) and **s** − **d** are both in day^−1^. Under mild diagonal dominance,

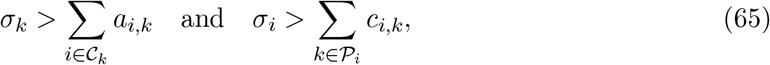

**M** is an *M* -matrix and **N**^∗^(**s**) = **A s** + **h** with **A** = **M**^−1^ ≥ 0 as before. The following generalizes Lemma 2.

#### Lemma 3

(Basal dominance on trophic DAGs). *Let* **v**^(*i*)^ = **Ae**_**i**_ *be the biomass response to a unit subsidy at level i. If all feeding links have efficiency κ <* 1 *and dissipation σ >* 0, *then for any upward trophic DAG with a unique basal node*,

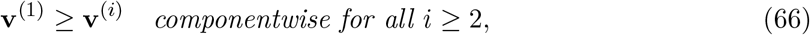

*with strict inequality for some component whenever any κ*_*j*_ *<* 1.

*Proof*. Each entry 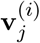 is a sum over directed paths from *i* to *j* of products of edge weights. Because every edge multiplies by a factor *<* 1 (conversion losses and dissipation), and because the basal node 1 has the shortest distance to all others, its influence vector **v**^(1)^ dominates the influence of any consumer subsidy. Strict inequality arises whenever some edge efficiency *κ <* 1 along a path forces attenuation. This extends the chain case to general DAGs.

#### Corollary 2

(Axis optimality under omnivory). *Under the same assumptions as Lemma 3, the optimizer of Q*(**s**) = **s**^*T*^**As** + **s**^*T*^**h** *with box constraints* 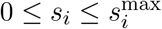 *still satisfiesss*

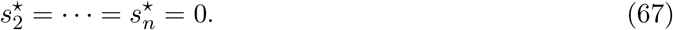

Thus axis optimality is robust to relaxing the strict chain assumption: direct consumer subsidies vanish at the entropy/output maximizer for any trophic DAG with *κ <* 1.

Omnivory modifies the numerical values of the bounds but not their form. Along the unsubsidized axis *s* = (*s*_1_, 0, …, 0),

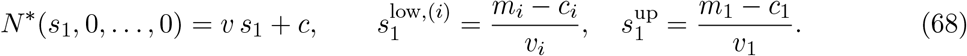

Adding additional prey routes increases *v*_*i*_ for *i* ≥ 2, thereby lowering consumer thresholds *s*^low^. At the same time, additional export reduces *v*_1_, thereby lowering the basal protection bound *s*^up^. Hence omnivory can either widen or shrink the emergence interval *I*_*L*_ = [max_*i*_ *s*^low,(*i*)^, *s*^up^]∩ [0, *s*^max^], depending on how energetic flows are redistributed.

Although omnivory can delay closure of the range by reducing consumer thresholds, the finite ceiling on trophic length still holds. With *κ <* 1 and *σ >* 0 bounded away from zero, attenuation of influence vectors up the DAG ensures *v*_*i*_ → 0 with trophic distance, so *s*^low,(*i*)^ → ∞ as *i* increases. Meanwhile *s*^up^ is finite and typically decreases as more consumers draw from the base. Thus the emergence interval *I*_*L*_ must close at finite *L*. Omnivory can shift the ceiling *L*^∗^ upward, but cannot remove it unless parameters approach thermodynamic limits (*κ* → 1, *σ* → 0).

The key results—axis optimality of the maximizer, the triad of bounds (*s*^low^, *s*^up^, *s*^max^), and the finite ceiling on trophic length—are therefore robust to structural generalizations from chains to trophic DAGs. Omnivory alters the numerical values of the bounds but not the qualitative mechanism: emergence intervals shrink and eventually vanish under cumulative energetic losses.

Additionally, our qualitative results extend to a broad class of consumer-resource nonlinearities (e.g., Holling II/III, Beddington-DeAngelis, ratio–dependent with saturation) under standard ecological conditions: (i) dissipative transfer and self–limitation (*σ*_*i*_ *>* 0); (ii) monotone, saturating gains; (iii) absence of positive feedback across levels so that the equilibrium Jacobian is Metzler with negative diagonal (a nonsingular M–matrix), which ensures an order–preserving equilibrium map *s* → *N*^∗^(*s*); and (iv) a rectangular supply cap 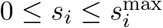. Under these assumptions, the basal–axis thresholds for a chain of length *L* are defined by solving the nonlinear steady state with 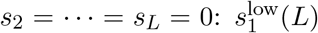and 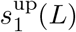 both increase with *L*, while 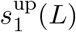 is clipped by 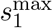 . Hence the emergence interval

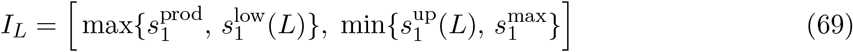

remains the correct object and collapses beyond a finite ceiling *L*^⋆^. Moreover, for 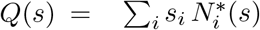, directional derivatives in higher–level subsidy directions are non–positive on the feasible set, yielding the same axis optimizer 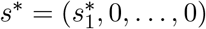.

